# Density-dependent selection at high food levels leads to the evolution of persistence but not constancy in *Drosophila melanogaster* populations

**DOI:** 10.1101/2022.06.05.494854

**Authors:** Neha Pandey, Amitabh Joshi

## Abstract

Mechanisms through which population dynamics evolve to be stable have been a subject of considerable interest in population biology. One of the ways through which population stability is likely to evolve is via density-dependent selection with or without an *r* and *K* trade-off. In this paper, we test whether the specific combination of egg number and food amount under which density-dependent selection is implemented affects the evolution of population stability attributes in *D. melanogaster* populations that have evolved under density-dependent selection for 75 generations. Our findings show that these populations have evolved higher persistence stability than controls, although constancy stability did not evolve. Moreover, these populations did not show an *r-K* trade-off, and evolved persistence largely through a significant decrease in sensitivity of growth rate to population density, especially at densities ranging from medium to the equilibrium population size. Qualitative comparison of these findings with those from another set of crowding-adapted *D. melanogaster* populations that had evolved both constancy and persistence stability, suggests that the ecology of larval crowding influences the consequent evolution of stability attributes. We discuss previous findings on the evolution of life-history traits to argue that differences in the ecology of density-dependent selection experienced at the larval stage affects population stability differently by altering the sensitivity of population growth rate to population density.

## INTRODUCTION

Following the demonstration that even simple, discrete-time, population growth models could show increasingly unstable and complex population dynamics with an increase in intrinsic population growth rate (*r*) (May 1974), it was thought that natural selection, all else being equal, would typically lead to higher intrinsic population growth rates and, therefore, to unstable dynamics. Yet, examination of population dynamics data from multiple species (Hassell *et al* 1976, Thomas *et al* 1980, Mueller and Ayala 1981 a) suggested that relatively stable population dynamics were quite common in nature (reviewed in Mueller *et al* 2000). This apparent contradiction between theoretical expectations and empirical data led to a growing interest in identifying evolutionary scenarios in which population would be likely to evolve to be stable. Early explanations for the evolution of population stability invoked group-selection for stable populations (Thomas *et al* 1980), as well as direct selection for reduced maximal or intrinsic growth rate (*r* in the logistic or Ricker models) (Hansen 1992, Ebenman *et al* 1996), A more plausible explanation was that of the evolution of population stability as a by-product of life-history evolution (Mueller and Ayala 1981 b), especially if the selected life-history-related traits happened to trade off with fecundity, thereby increasing population stability (Turelli and Petry 1980, Stokes *et al* 1988, Gatto 1993, Ebenman *et al* 1996, Prasad *et al* 2003).

Under density-dependent selection, population evolving at low density are expected to evolve high intrinsic growth rate (*r*), while a higher population size at equilibrium (*K*) is expected to evolve under high population density (MacArthur and Wilson 1967). While testing the hypotheses from density-dependent selection theory, Mueller and Ayala (1981 b) subjected *Drosophila* populations to low density (*r*-type) and high density (*K*-type) environments and observed evolved differences in the growth rates at low and high density and trade-offs between them. Following these observations of the effects of density on traits that influenced *r* and *K*, Mueller and Ayala (1981 b) proposed that population dynamics could evolve to be more stable if selection in chronically crowded conditions led to the evolution of traits that lowered *r* as a correlated response to selection for traits that increased *K*.

The possible role of density-dependent selection in mediating the evolution of population stability was first examined in *D. melanogaster* populations which were specifically selected for adaptations to crowding experienced at the larval stage (Mueller *et al* 2000). These CU populations faced crowding as larvae but were uncrowded at the adult stage, while their ancestral controls (UU) populations did not experience crowding in either larval or adult stage. Selection under high larval crowding in the CUs led to the evolution of traits very similar to those seen earlier in the *D*. *melanogaster* populations used by Mueller and Ayala (1981 b), most notably the evolution of increased larval feeding rates at the cost of efficiency of food conversion to biomass (Joshi and Mueller 1996), but an examination of their population dynamics did not show any evolved differences in constancy stability (sensu Grimm and Wissel 1997); in that study very large populations were used and no extinctions were observed (Mueller *et al* 2000, Mueller and Joshi 2000).

Subsequently, support for the evolution of population stability through density-dependent selection was found in *D. ananassae* populations selected for adaptation to larval crowding (ACUs: Dey *et al* 2012), which showed evolutionary increase in both constancy and persistence stability (*sensu* Grimm and Wissel 1997) as compared to their uncrowded controls (ABs). This study also suggested that the evolution of greater population stability in the ACUs was partly mediated through an *r*-*K* trade-off. It is worth noting that the ACU populations had evolved increased competitive ability through greater time efficiency of food to biomass conversion, without evolution of increased larval feeding rate (Nagarajan *et al* 2016), as opposed to the crowding-adapted CU populations of Mueller *et al* (2000). These differences in which traits evolved under larval crowding were finally attributed to the different combination of egg number and food amount at which the ACU and CU populations, respectively, experienced chronic larval crowding, with the ACUs being selected at very low food amounts (Nagarajan *et al* 2016, Sarangi *et al* 2016).

Based on the differences in stability evolution between the CUs and ACUs, Dey *et al* (2012) speculated that the specific traits that evolve in response to larval crowding, and whether they mostly affect the effectiveness or tolerance components of competitive ability (sensu Joshi *et al* 2001), could possibly help determine whether or not population stability evolved as a correlated response to density-dependent selection. To further test this idea, we studied two different sets of crowding-adapted *D*. *melanogaster* populations that shared common ancestry with the CU populations of Mueller *et al* (2000). One set of populations (MCUs) experienced chronic larval crowding at the same combination of egg number and food amount as the ACU populations. Another set of populations (LCU) were subjected to chronic larval crowding under egg number and food amount combination approximating that used for the CUs The derivation and maintenance of the MCU and LCU populations is described in detail by Sarangi (2018). When we compared the MCUs and their controls for population stability, we found that, similar to the ACUs, the MCUs had evolved greater constancy and persistence stability, but without the involvement of an *r*-*K* trade-off (Pandey and Joshi 2022). Here, we examine population stability in the LCUs and their controls (the same controls as the MCUs), specifically asking whether they show results similar to the CUs of Mueller *et al* (2000), especially since the LCUs are known to have evolved higher larval feeding rates (Sarangi 2018), like the CUs but not the MCUs. Any observed differences between how population dynamics and stability characteristics have evolved in the LCUs as compared to the MCUs would permit a rigorous experimental test of the speculative predictions of Dey *et al* (2012) about how the specific egg number and food amount combination at which larval crowding is experienced can affect whether or not population stability evolves.

## MATERIALS AND METHODS

### Experimental populations

We used eight large outbred lab-maintained populations (four selected and four controls) of *D. melanogaster* which are maintained on a 21-day discrete generation cycle in constant light at 25°C±1°C with around 80 percent humidity: four **LCU** populations (<u>L</u>arry Mueller CU-type, <u>C</u>rowded as larvae and <u>U</u>ncrowded as adults), and four **MB** (<u>M</u>elanogaster <u>B</u>aseline, serves as ancestral controls) populations (complete details are given in Sarangi 2018). LCUs are subjected to competition at larval stage at relatively high food amounts (hence, selected for adaptation to larval crowding), while MBs do not face larval competition for food. The LCUs are maintained in 6-dram glass vials (9 cm height *×* 2-2.2 cm inner diameter) at a density of `∼`1200 eggs per 6 mL corn meal food, and at the adult stage at `∼`1800 adults in Plexiglas cages (dimension 25 *×* 20 *×* 15 cm^3^). The eclosing flies from LCU culture vials are transferred to their respective cages every day after the 8^th^ day from egg collection till day 20 post egg lay. In the cages, these flies are given corn meal food change every alternate day in a Petridish, and the cage contains a moist cotton ball which is changed every alternate food change. On the 18^th^ day post egg lay, a Petridish containing a generous amount live acetic acid yeast paste is provided for `∼`2.5 days after which a cut plate (vertical food surface) is provided (on the 20^th^ day post egg lay) for females to lay eggs for `∼`18 hours, after which (on the 21^st^ day) eggs are roughly counted to 1200 eggs and placed in vials containing 6 mL of food. The MBs are maintained similarly to the LCUs, except that they are collected at an egg density of `∼`70 eggs/ ∼6 mL food in 8-dram vials (9.5 cm height *×* 2.2-2.4 cm inner diameter). Also, all eclosing flies from MB populations are collected at once on the 11^th^ day after egg collection into cages, as most adults eclose by then in the absence of larval competition. Each MB population consists of 40 vials at the larval stage, as opposed to 12 vials for each LCU population, in order to maintain similar adult density (`∼`1800 adults).

### Population dynamics experiment

We carried out the population dynamics experiment for 26 generations in a destabilizing LH food regime (L = low quantity of larval food, and H = high quantity of adult food with yeast supplement: Mueller and Huynh 1994, Sheeba and Joshi 1998). This experiment was carried out with 1 mL of larval food in the LH environment, as this food regime provides a high probability of being able to detect differences in constancy and persistence (Vaidya 2013, Pandey and Joshi 2022). At the time of initiating the population dynamics experiment, the LCU populations had undergone about 75 generations of selection and had diverged from their controls in many traits relevant to fitness under larval crowding (Sarangi 2018).

We started the experiment by deriving 10 single-vial populations from each of the eight LCU and MB populations, after one generation of common rearing at low larval density to eliminate any non-genetic parental effects. We started each vial population with 8 mated females which were allowed to lay eggs in in the vial for 24 hours (counted as 16 adults in generation 0). We began transferring eclosing flies from egg vials to matched adult collection vials containing around 4 mL of cornmeal food, after day 8 from egg lay. As eclosion is spread out over several days due to larval competition, we transferred eclosing flies to their respective adult collection vials every day, till day 18 post egg lay. Correspondence between egg vials and adult collection vials was meticulously maintained, and all fly-transfers were done with extreme care to avoid losing any flies. We shifted adults to fresh adult collection vials every alternate till day 18 post egg lay. On day 18, provided flies with a dab of live acetic-acid yeast on the wall of a fresh adult collection vial to boost adult fecundity. On day 20 from egg collection, we transferred flies from the adult vials into new egg-laying vials with 1 mL food and allowed them to lay eggs over the next 16 hours to lay eggs for next-generation. The adults were then moved into empty vials for the census counts after freezing. Any fly found dead during the 16-hour egg-laying phase was also included in the census count.

The eggs laid by the flies in each vial initiated the next generation, i.e. density was not controlled. In parallel with the vials described above, we maintained a set of five backup vials per population whose maintenance was similar to the experimental vial populations except that backup vial populations were maintained at a low larval density. Each generation, we randomly chose 5 females from each backup vial population to lay eggs for 16 hours in 6 mL of food to start the next backup generation, while the rest of the flies were discarded. Following Dey and Joshi (2006), we maintained these backup vials to reset the experimental populations (with 4 males and 4 females) in case of extinction (absence of even one male-female pair) in a vial population on day 20 post egg lay. A total of 80 single-vial populations (2 selection regimes × 4 replicate populations × 10 single-vial populations) were, thus, censused over the 26 generation long population dynamics experiment. These 80 time series of population size data, along with the number of times each population went extinct over the 26 generations, constituted the primary data for further analyses.

### Stability indices

#### Constancy

We compared constancy stability (*sensu* Grimm and Wissel 1997) in MBs and LCUs using two indices: coefficient of variation (CV) in population size, and fluctuation index of population size (FI). Coefficient of variation (CV) in population size reflects dispersion, scaled by the mean, around the mean population size. We also assessed constancy through FI which measures the mean one-step absolute change in population size, scaled by the mean population size (Dey and Joshi 2006), as

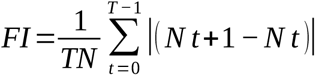

where *T* is the number of generations, *N* is the average population size, and *N_t_* and *N_t_*_+1_ are the population sizes at generations *t* and *t*+1, respectively. Constancy was interpreted as being the inverse of CV or FI, respectively.

#### Persistence

We compared persistence stability between MBs and LCUs using the frequency of extinction per generation in each single-vial population. Persistence was interpreted as the inverse of the extinction probability. We counted consecutive extinctions in the same population as one extinction because consecutive extinctions in experiments like these are typically not independent (Dey *et al* 2008). We also calculated mean population size of each single-vial population across the 26 generations of the experiment.

### Demographic attributes

In all the single-vial populations, we examined three demographic attributes: intrinsic population growth rate, equilibrium population size, and sensitivity of realized population growth rate to population density. It is known that the dynamics of single-vial *Drosophila* populations in the LH food regime are captured reasonably well by the Ricker (1954) model (Sheeba and Joshi 1998). However, the responses of *Drosophila* population dynamics to various food or selection regimes need not necessarily be limited by any simple population growth model (Tung *et al* 2019, Joshi 2022). Consequently, we examined these attributes in different ways, some based on the Ricker model and others directly based on the empirical data.

In the Ricker-based approach, we estimated *r* and *K*, representing intrinsic population growth rate and equilibrium population size, respectively, as well as *α* = *r*/*K*, reflecting the sensitivity of realized population growth rate to density. These estimations involved either (a) plotting a regression line between Ln (*N_t_*_+1_/*N_t_*) on the Y-axis and *N_t_* on the X-axis, and taking the Y-intercept, X-intercept and slope as estimates of *r*, *K* and *α*, respectively, or (b) using non-linear curve fitting (following Dey *et al* 2008), through the Quasi-Newton method (StatSoft 1995), followed by calculating *α* as -*r*/*K*.

We also examined realized population growth rates (*N_t_*_+1_/*N_t_*) at low (*N_t_* ≤ 30) and high (*N_t_* > 60) densities, as correlates of *r* and *K*, respectively, following the logic of Joshi *et al* (2001). We also checked the realized population growth rates at different cut-off values for low and high density to assess the robustness of the result. Finally, we estimated realized population growth rates (*N_t_*_+1_/*N_t_*) over the entire range of population densities observed in the single-vial populations, in bin sizes of 30.

### Comparison between LCUs and MCUs

Although, our purpose in this study was to see whether populations adapted to chronic larval crowding at high versus low food amounts differed in the demographic and stability characteristics they evolved due to density-dependent selection, it was not possible to directly compare evolutionary change in the LCUs and MCUs since, for logistical reasons, the two population dynamics experiments could not be run together. Therefore, we compared them indirectly, making use of the fact that both the LCUs and the MCUs were derived from the same four ancestral control (MB) populations.

We used data from the study in this chapter, and the one described in Pandey and Joshi 2022, and transformed the estimated values of stability indices, mean population sizes and the three Ricker-based demographic attributes (*r*, *K* and *α*) into fractional deviations from control population values. For each measure from each single-vial population in the LCU and MCU population dynamics experiments, we calculated *Y*\**_ij_* = (*Y_ij_* – *µ_j_*) / *µj* (*i* = 1…10, *j* = 1…4), where *Y*\**_ij_* was the transformed response variable, *Y_ij_* was the measure of a given attribute in the *i*^th^ replicate single-vial population (in the population dynamics experiment) of the *j*^th^ replicate population (from the ongoing selection experiment), and *µ_j_* was the mean value of that attribute in the *j*^th^ replicate population of the control MBs, averaged over all 10 single-vial populations within that replicate. These transformed response variables were subsequently used as input data for further analyses.

### Statistical analyses

To compare the population dynamics and stability characteristics of the crowding-adapted LCUs with their controls (MBs), we used mixed model analysis of variance (ANOVA) to analyze all the response variables. The ANOVA models included selection regime as a fixed factor with two levels, crossed with random blocks (four levels) representing the common ancestry of LCU-*i* and MB-*i*. and food level, We performed separate ANOVAs on the coefficient of variation in population size, fluctuation index, extinction probability, average population size, intrinsic growth rate (estimated through three different methods), equilibrium population size (estimated through three different methods), and the sensitivity of realized population growth rate to population density (estimated through two methods). The same ANOVA design was used for the LCU-MCU comparison, using the transformed response variables (see precding sub-section): here, the two levels of selection regime were LCU and MCU, rather than LCU and MB, A separate ANOVA was performed on realized growth rates corresponding to different population size bins, with bin as an additional fixed factor, crossed with selection regime and block. All analyses were performed in Statistica Ver. 5.0 (StatSoft 1995), and post-hoc comparisons used Tukey’s HSD test at *P* = 0.05.

## RESULTS

### Constancy and persistence stability

We found that constancy stability was not significantly different between the crowding-adapted LCUs and their controls, the MBs, using either the CV in population size or the fluctuation index (FI) (Fig. 1, Table 1). Indeed, both CV and FI hardly differed on average between the LCU and MB populations (Fig. 1), clearly indicating that constancy stability has not evolved in the LCU populations.

**Figure 1.**
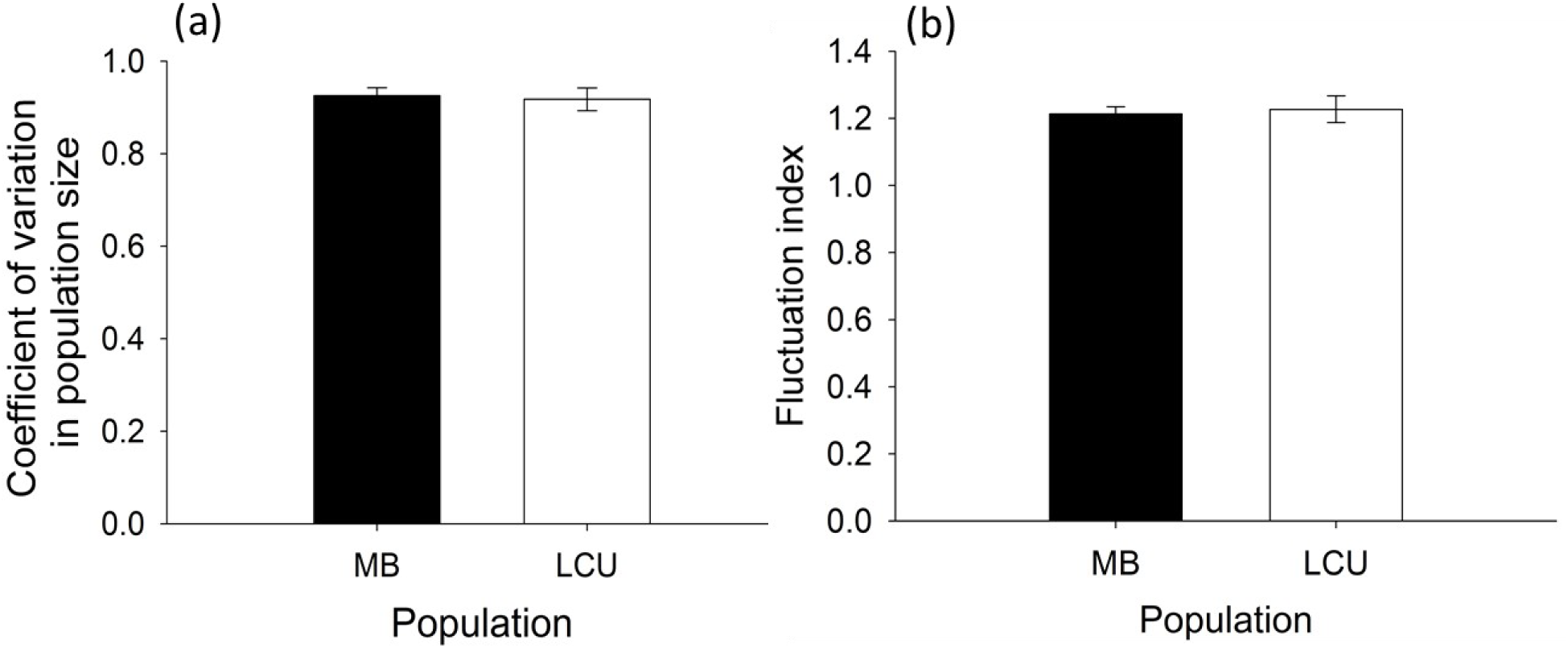
Constancy stability in MB and LCU populations: (a) mean coefficient of variation in population size, and (b) mean fluctuation index. Error bars around the means are standard errors based on variation among the means of the four replicate populations within each selection regime.

**Table 1:**
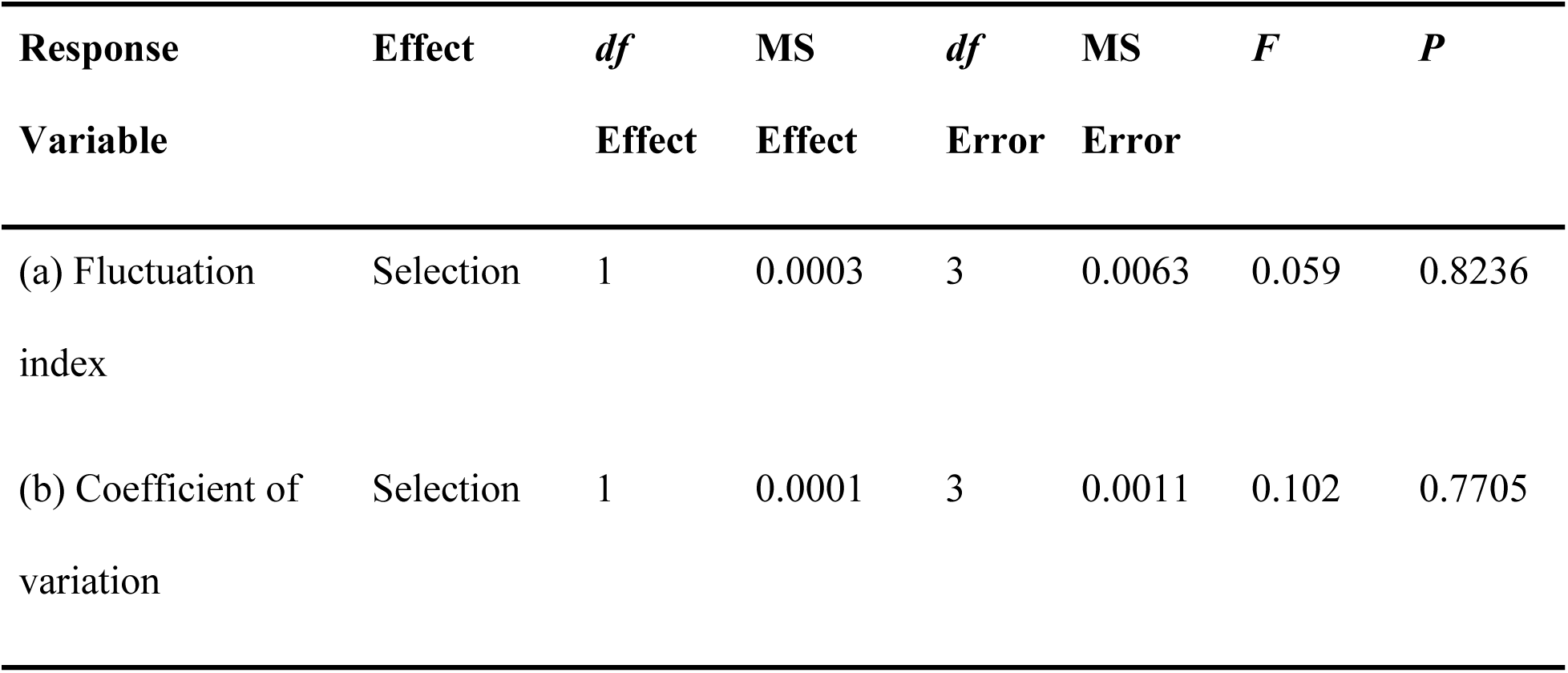
Summary results of ANOVA done on constancy stability measured as (a) coefficient of variation in population size, and (b) fluctuation index. The table shows the main effect of selection (LCUs vs MBs). Since we were primarily interested in fixed main effects, block effects and interactions have been omitted for brevity.

| **Response** | **Effect** | ***df*** | **MS** | ***df*** | **MS** | ***F*** | ***P*** |
| --- | --- | --- | --- | --- | --- | --- | --- |
| **Variable** |  | **Effect** | **Effect** | **Error** | **Error** |  |  |
| (a) Fluctuation index | Selection | 1 | 0.0003 | 3 | 0.0063 | 0.059 | 0.8236 |
| (b) Coefficient of variation | Selection | 1 | 0.0001 | 3 | 0.0011 | 0.102 | 0.7705 |

In contrast, greaterpersistence stability has evolved in the LCUs, as their extinction rate was signficantly lower than the MBs (Fig. 2 a, Table 2 a), with LCU populations being nearly half less likely to go extinct as compared to MB populations.

**Figure 2.**
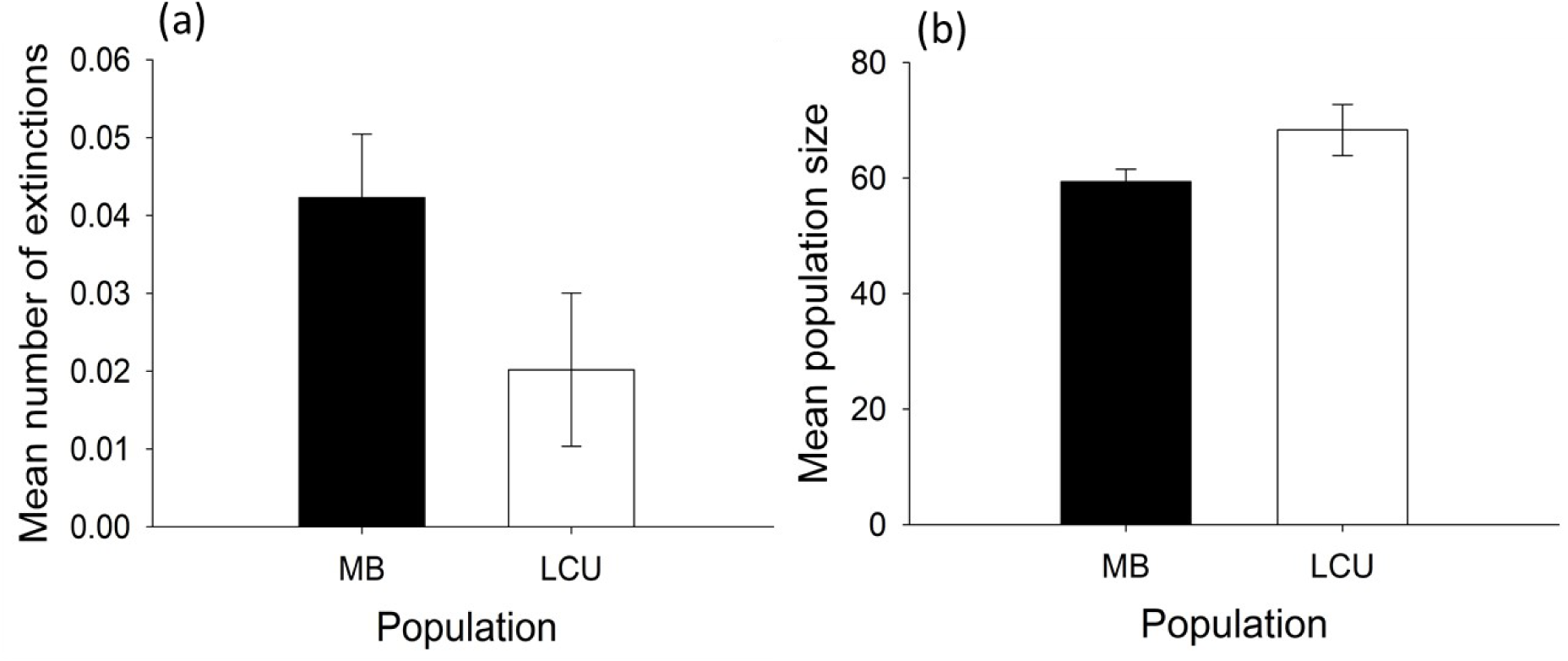
Persistence stability and average population size for MB and LCU populations: (a) mean number of extinctions per generation, and (b) mean population size. Error bars around the means are standard errors based on variation among the means of the four replicate populations within each selection regime.

**Table 2:**
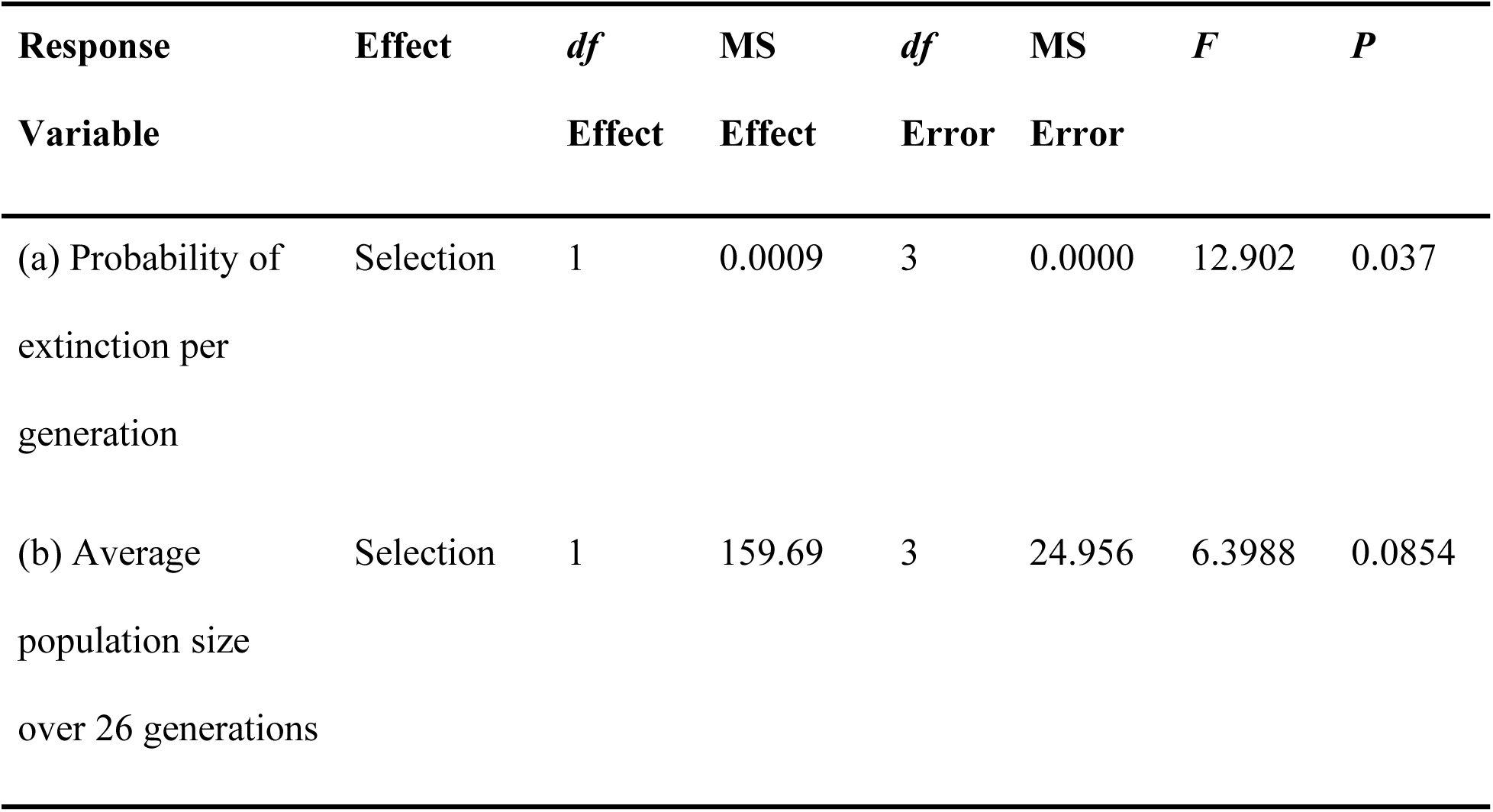
Summary results of ANOVA done on (a) persistence stability measured as probability of extinction, and (b) average population size. Since we were primarily interested in fixed main effects, block effects and interactions have been omitted for brevity.

### Mean population size

We found that although the LCUs showing slightly higher mean population size as compared to MBs (Fig. 2 b), the difference was not statistically significant (Table 2 b).

### Ricker-based demographic attributes

Intrinsic rate of population growth (*r*), whether estimated from linear regression of Ln (*N_t+_*_1_/*N_t_*) on *N_t_* , or via non-linear fitting, did not differ significantly between MBs and LCUs (Table 3 a, b). Not only were differences in *r* between LCUs and MBs not significant, even the magnitude of the differences was negligible (Fig. 3 a, b)

**Figure 3.**
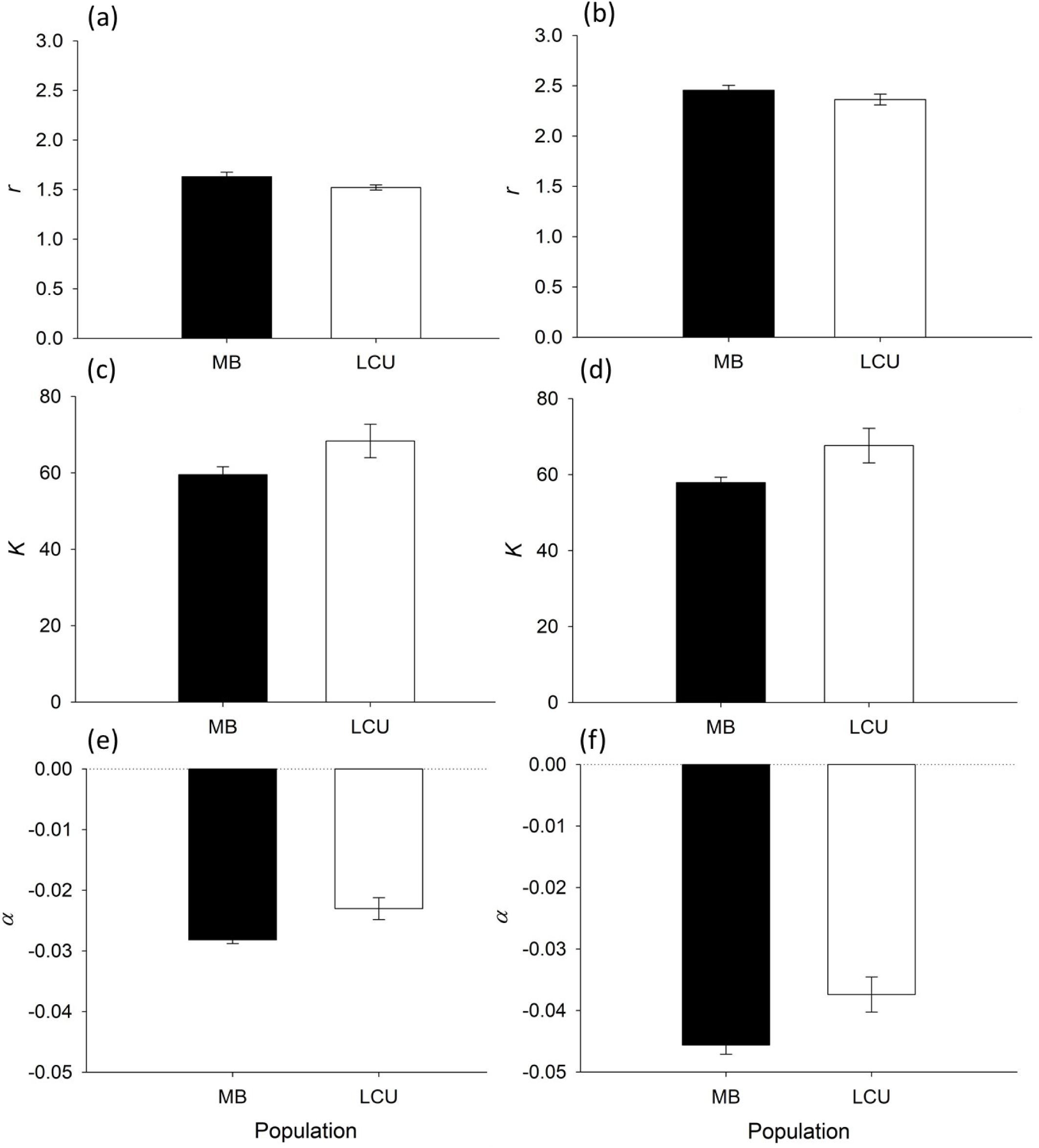
Mean demographic attributes of MB and LCU populations, based on the Ricker equation: (a) intrinsic growth rate *r* (estimated by taking the Y-intercept of the regression line between Ln (*N_t+_*_1_/*N_t_*) on Y-axis and *N_t_* on X-axis), (b) intrinsic growth rate *r* (estimated by non-linear fitting), (c) equilibrium population size *K* (estimated by taking the X-intercept of the regression line between Ln (*N_t+_*_1_/*N_t_*) on Y-axis and *N_t_* on X-axis), (d) equilibrium population size *K* (estimated by non-linear fitting), (e) sensitivity of realized growth rate to density *α* (estimated by taking the slope of the regression line between Ln (*N_t+_*_1_/*N_t_*) on Y-axis and *N_t_* on X-axis), and (f) sensitivity of realized growth rate to density *α* (estimated by non-linear fitting). Error bars around the means are standard errors based on variation among the means of the four replicate populations within each selection regime.

**Table 3:** Summary results of ANOVA done on the Ricker-based estimates of intrinsic population growth rate (*r*), equilibrium population size (*K*), and sensitivity of realized population growth rate to population density (*α*), estimated from either linear regression of Ln (*N_t_*_+1_/*N_t_*) on *N_t_* (a, c, e), or non-linear curve fitting (b, d, f). Since we were primarily interested in fixed main effects, block effects and interactions have been omitted for brevity.

| **Response** | **Effect** | ***df*** | **MS** | ***df*** | **MS** | ***F*** | ***P*** |
| --- | --- | --- | --- | --- | --- | --- | --- |
| **Variable** |  | **Effect** | **Effect** | **Error** | **Error** |  |  |
| (a) $r$ (Linear regression) | Selection | 1 | 0.0239 | 3 | 0.0052 | 4.591 | 0.1215 |
| (b) $r$ (Non-linear fitting) | Selection | 1 | 0.0168 | 3 | 0.0190 | 0.89 | 0.4157 |
| (c) $K$ (Linear regression) | Selection | 1 | 155.5 | 3 | 25.435 | 6.113 | 0.0898 |
| (d) $K$ (Non-linear fitting) | Selection | 1 | 189.89 | 3 | 27.454 | 6.916 | 0.078 |
| (e) $\alpha$ (Linear regression) | Selection | 1 | 0.000 | 3 | 0.000 | 15.382 | 0.029 |
| (f) $\alpha$ (Non-linear fitting) | Selection | 1 | 0.0001 | 3 | 0.000 | 16.273 | 0.0273 |

The equilibrium population size (*K*) of LCUs was somewhat higher than the MBs for both methods of estimation: linear regression of Ln (*N_t+_*_1_/*N_t_*) on *N_t_* and non-linear fitting (Table 3 c, d), but the differences were not significant (Fig. 3 c, d).

The sensitivity of realized population growth rate to population density (*α*) showed that LCUs were significantly less sensitive to change in population density than MBs, regardless of the method of estimation (Fig. 3 e, f, Table 3 e, f).

### Model-free demographic attributes

When we compared empirically estimated mean realized population growth rates at low versus high density in the LCUs and MBs, the only significant ANOVA effect was that of density (Table 4). Essentially the magnitude of the difference between growth rates at low versus high density rendered the effect of all other sources of variation on realized population growth rate relatively negligible (Fig. 4 a).

**Figure 4.**
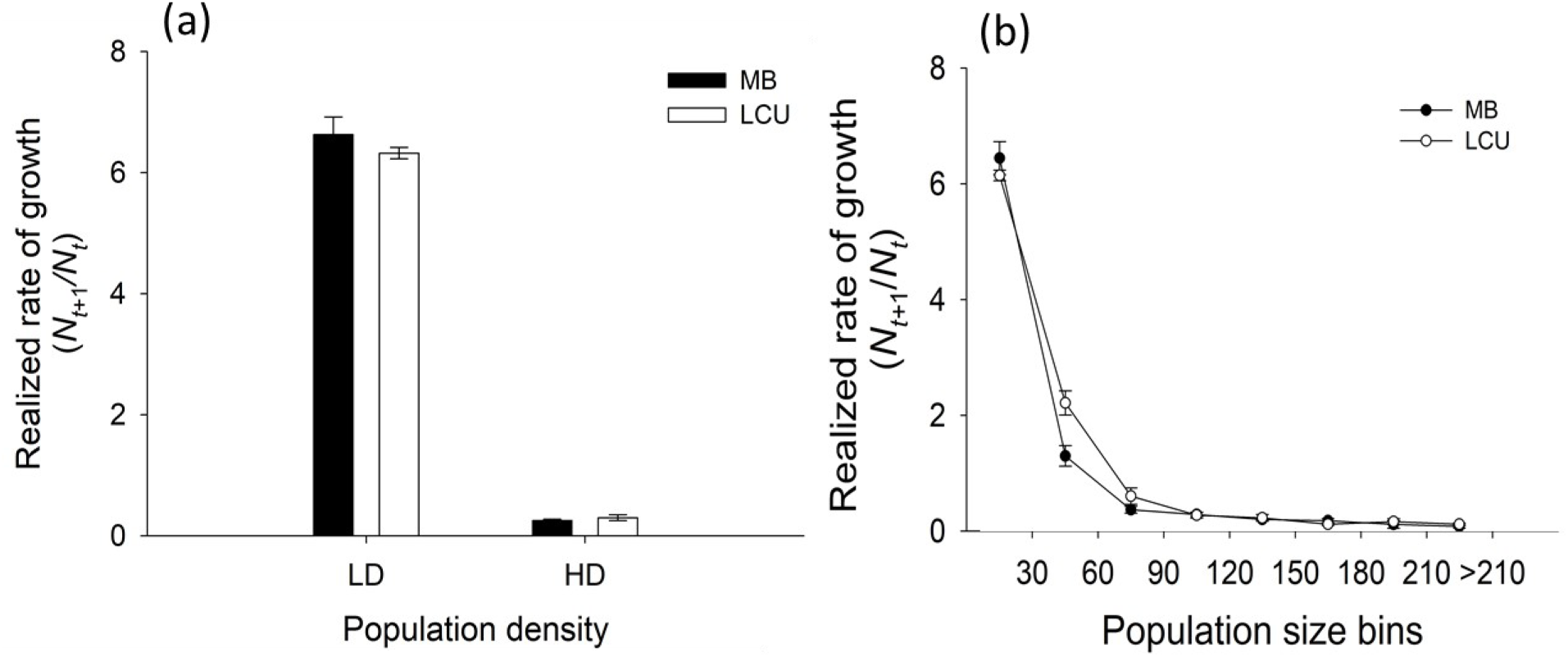
Mean empirical realized growth rates (*N_t+_*_1_/*N_t_*) of MB and LCU populations: (a) at low ( *N_t_* < 30) and high (*N_t_* > 60) density, and (b) across the range of densities seen in the single-vial populations, in bins of 30. population bin sizes of 30. Error bars around the means are standard errors based on variation among the means of the four replicate populations within each selection regime.

**Table 4:**
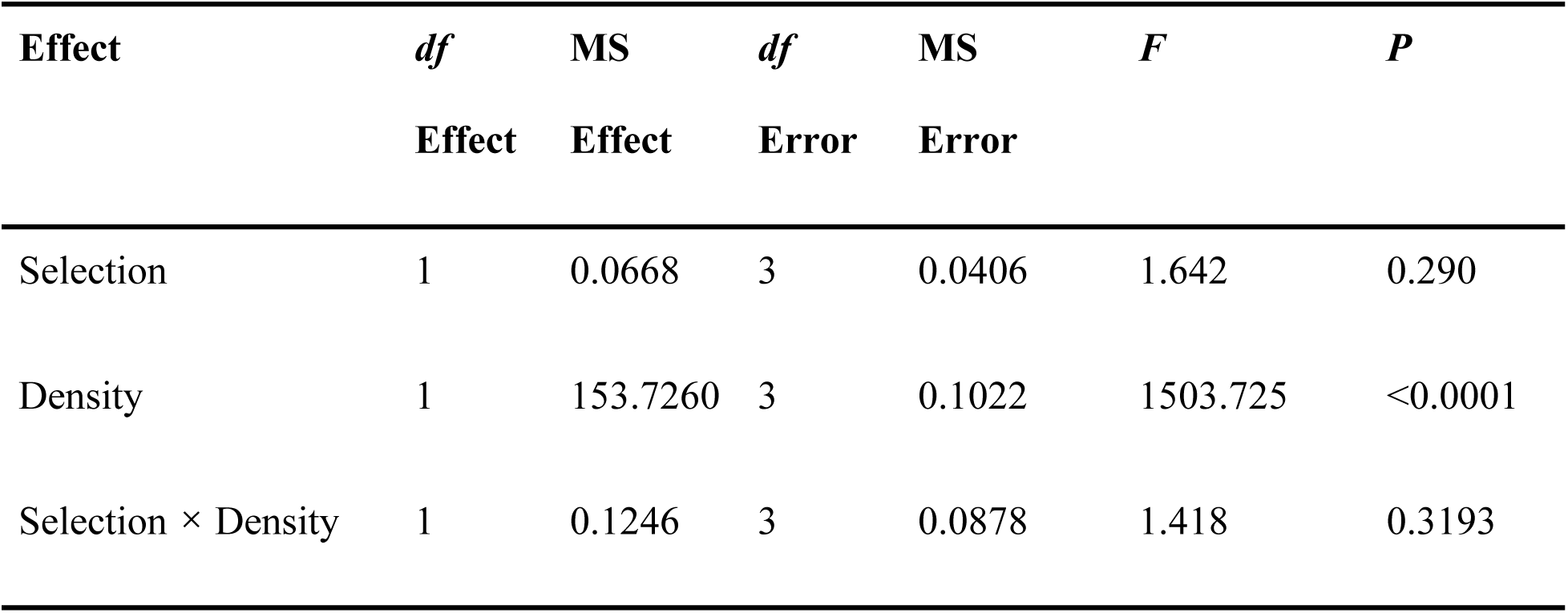
Summary results of ANOVA done on realized growth rate (*N_t_*_+1_/*N_t_*) in MBs and LCUs at low and high population densities. The table shows the main effect of selection, population density and their interaction. Since we were primarily interested in fixed main effects and interactions, block effects and interactions have been omitted for brevity.

The picture became slightly clearer when we examined mean realized population growth rate across the full range of densities achieved in the single-vial populations, in bin sizes of 30 (Fig. 4 b). There were significant ANOVA effects of population size bin (density level), and the interaction between selection regime and population size bin, for mean realized population growth rate data (Table 5) food. On an average, realized population growth rates were higher at lower population densities until a reasonably high density was attained (*N_t_* > 75), beyond which point realized population growth rates tended to level off (Fig. 4 b). The significant interaction between selection regime and population size bin was driven by the fact that LCUs sustained significantly higher mean realized population growth rate than MBs at densities between 30 and 60 individuals per vial; differences between LCUs and MBs at other bins were not significant in the post-hoc comparisons (Fig. 4 b).

**Table 5:** Summary results of ANOVA done on realized population growth rate (*N_t+_*_1_/*N_t_*) at different population density (*N_t_*) bins in MBs and LCUs with a bin size of 30. Since we were primarily interested in fixed main effects and interactions, block effects and interactions have been omitted for brevity.

| **Effect** | ***df*** | **MS** | ***df*** | **MS** | ***F*** | ***P*** |
| --- | --- | --- | --- | --- | --- | --- |
|  | **Effect** | **Effect** | **Error** | **Error** |  |  |
| Selection | 1 | 0.2084 | 3 | 0.0292 | 7.1161 | 0.0758 |
| Bin | 7 | 38.2231 | 21 | 0.0452 | 845.5378 | <0.0001 |
| Selection × Bin | 7 | 0.2657 | 21 | 0.0537 | 4.9421 | 0.0019 |

### Differences in the stability and demographic attributes of MCUs and LCUs

After transformation of various response variables pertinent to population dynamics and stability in the LCUs and MCUs, expressing their values as a fractional difference from the MB controls in the respective population dynamics experiments, ANOVAs revealed a significant difference between MCUs and LCUs only in their constancy stability as reflected by CV in population size (Table 6). For all other response variables, differences between MCUs and LCUs were not significant (Table 6).

**Table 6.** Summary results of ANOVA done on the scaled differences of MCUs and LCUs from their common controls (MBs) for various stability- and dynamics-related response variables. The table shows the main effect of population type (LCUs or MCUs). Since we were primarily interested in fixed main effects and interactions, block effects and interactions have been omitted for brevity.

| **Response** | **Effect** | ***df*** | **MS** | ***df*** | **MS** | ***F*** | ***P*** |
| --- | --- | --- | --- | --- | --- | --- | --- |
| **Variable** |  | **Effect** | **Effect** | **Error** | **Error** |  |  |
| (a) Fluctuation index | Population type | 1 | 0.0075 | 3 | 0.0055 | 1.358 | 0.328 |
| (b) Coefficient of variation | Population type | 1 | 0.0212 | 3 | 0.002 | 10.464 | 0.048 |
| (c) $r$ (Linear regression) | Population type | 1 | 0.0266 | 3 | 0.0041 | 6.365 | 0.0859 |
| (d) $r$ (Non-linear fitting) | Population type | 1 | 0.0002 | 3 | 0.0052 | 0.0430 | 0.8489 |
| (e) $K$ (Linear regression) | Population type | 1 | 0.0702 | 3 | 0.0174 | 4.0176 | 0.1387 |
| (f) $K$ (Non-linear fitting) | Population type | 1 | 0.0673 | 3 | 0.0163 | 4.1145 | 0.1355 |
| (g) $\alpha$ (Linear regression) | Population type | 1 | 0.0029 | 3 | 0.0088 | 0.3322 | 0.6047 |
| (h) $\alpha$ (Non-linear fitting) | Population type | 1 | 0.0318 | 3 | 0.0046 | 6.849 | 0.0792 |

## DISCUSSION

Our results essentially confirmed the insight of Dey *et al* (2012) that *Drosophila* populations adapting to chronic larval crowding at high versus low food amounts are likely to differ in whether or not they also evolve greater population stability attributes as a correlated response to density-dependent selection. In terms of demographic attributes and population stability characteristics, we found that the LCUs, adapted to larval crowding at relatively high food amounts, had evolved a different pattern of responses relative to controls than the MCUs (Pandey and Joshi 2022), which were adapted to larval crowding at very low food amounts. The LCUs did not evolve higher constancy than controls (Fig. 1, Table 1), but did evolve higher persistence stability (Fig. 2 a, Table 2 a), presumably largely through the evolution of lower sensitivity of growth rate to density (less negative *α*: Fig. 3 e, f, Table 3 e, f) and, perhaps, a slight tendency, though not significant, towards higher *K* (Fig. 3 c, d, Table 3 c, d) and average population size (Fig. 2 b, Table 2 b) than the MB controls. The evolution of persistence but not constancy in the LCUs also supports the previous view (Dey *et al* 2008) that these two stability attributes do not necessarily coevolve, although they can in some circumstances (e.g. Dey *et al* 2012, Pandey and Joshi 2022). The empirical estimates of realized population growth rates in the LCUs across densities also revealed a difference from what was seen in the case of the MCUs (Pandey and Joshi 2022). The MCUs exhibited elevated realized population growth rates, compared to MB controls, across a wide range to medium to high densities, spanning both below and above the equilibrium populations size (Fig. 5 in Pandey and Joshi 2022). As noted by Pandey and Joshi (2022), this kind of change cannot be accommodated within the framework of even the *θ*-Ricker or *θ*-logistic models. The LCUs, on the other hand, showed higher realized population growth rates than MBs across a narrower range of medium to high densities, mostly spanning densities less than or up to the equilibrium population size (Fig. 4 b). This pattern of evolution could perhaps be explainable, in principle, by different degrees to which the sensitivity of various fitness components to density has evolved in the MCUs versus the LCUs, and is something that needs to be investigated further. We also note that the kind of change in realized population growth rates at medium to high densities below *K* seen in the LCUs can be modeled via evolutionary change in *θ* using models like the *θ*-Ricker or *θ*-logistic.

We discuss these results in the context of the mechanisms through which population stability can evolve, especially through changes in the pattern of density-specific realized population growth rates, reflected in parameters like *r*, *K* and *α* in simple population growth models like the logistic or Ricker. We also discuss how ecological differences in density-dependent selection at the larval stage might influence the evolution of population stability by comparing various fitness-related traits, and the population dynamics and stability attributes of the LCUs with other populations that have experienced larval crowding at various combinations of egg density and food volume than the LCUs (Table 7).

**Table 7.**
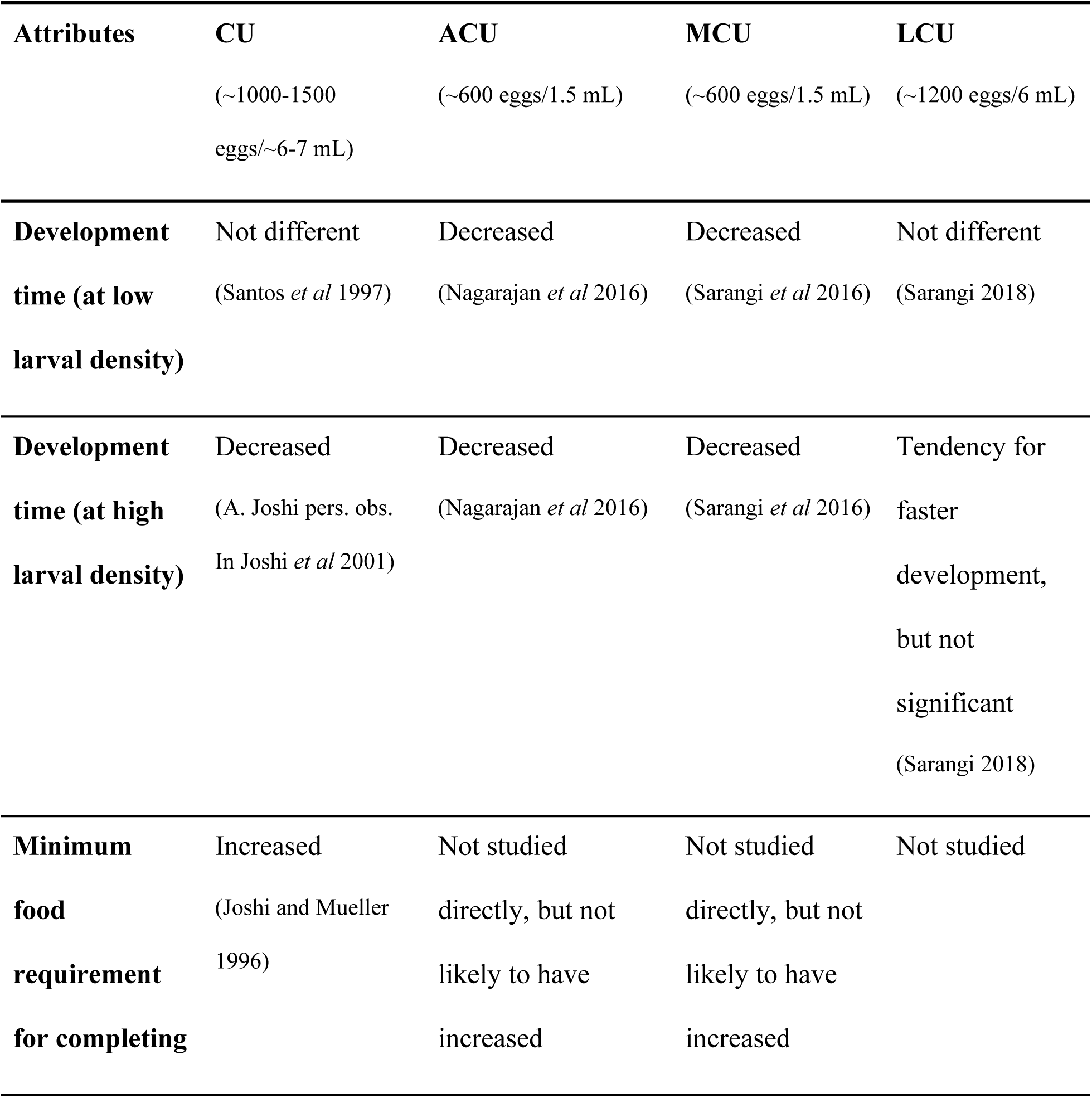
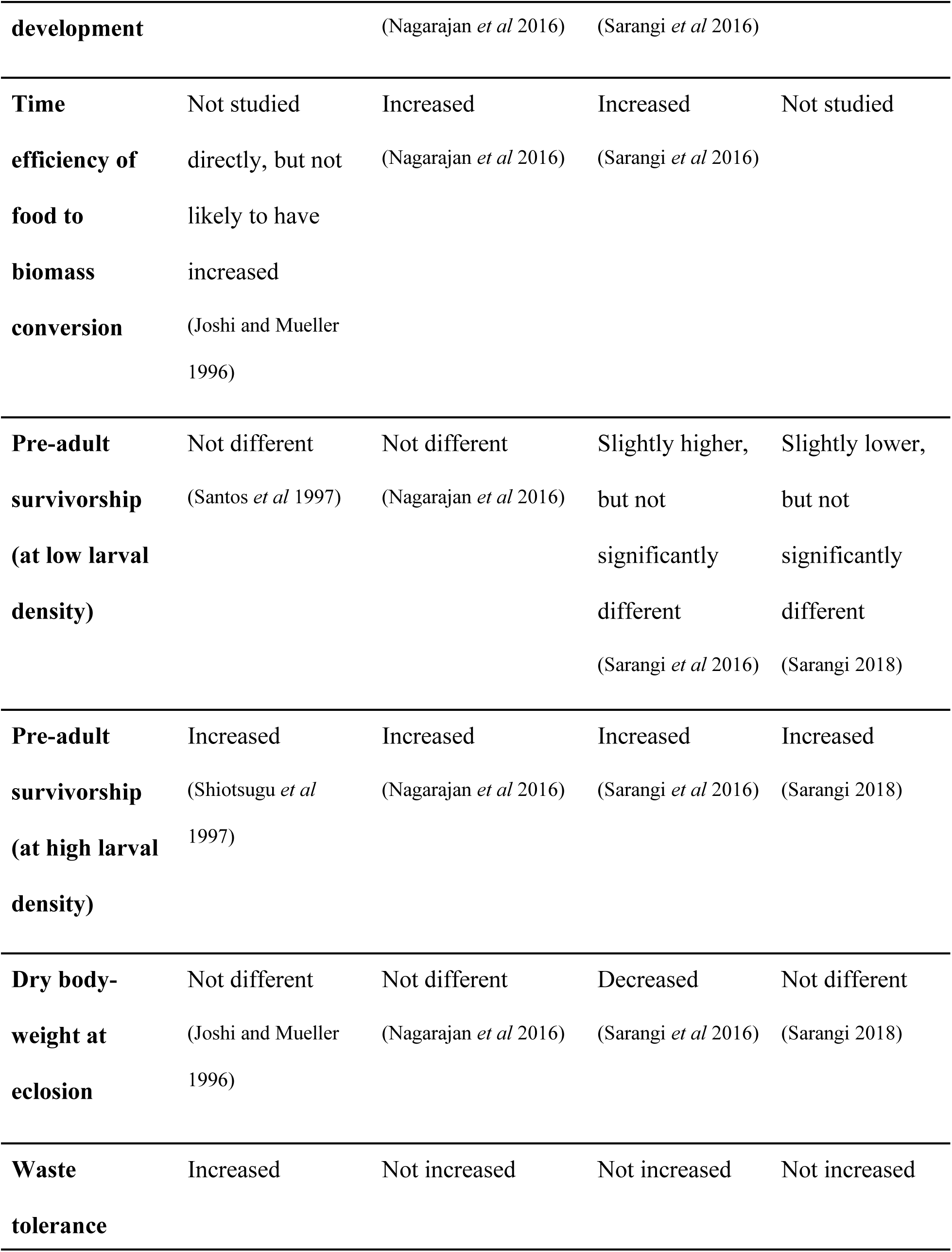
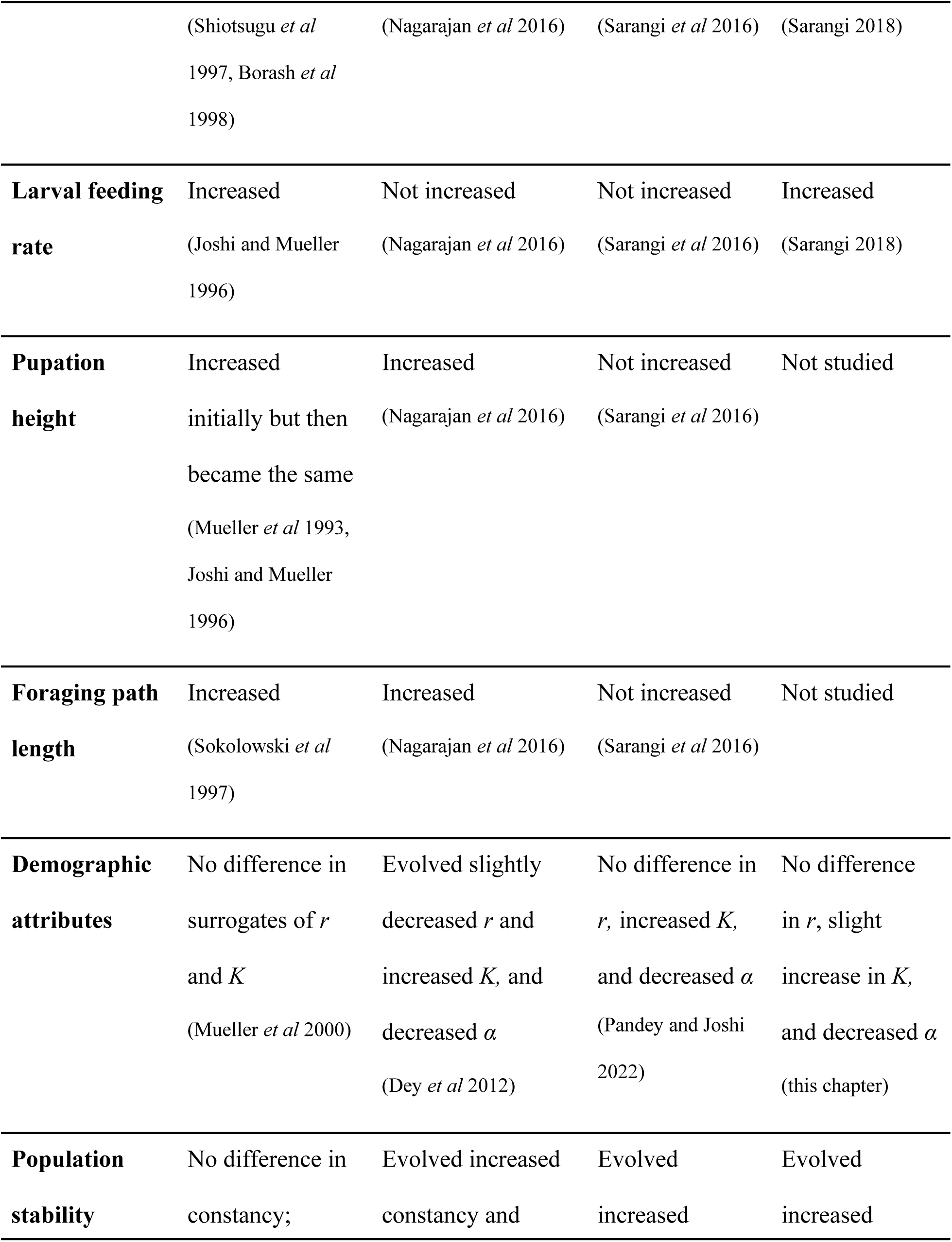
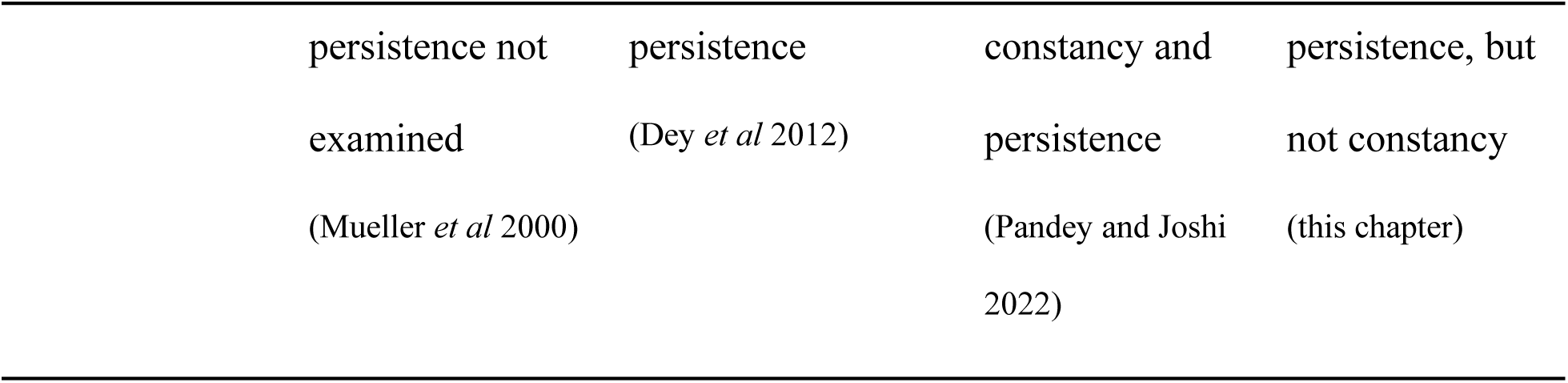
Comparison of various traits relevant to fitness at high larval density, demographic attributes and population stability characteristics, in the various *Drosophila* populations selected under density-dependent selection that have been investigated for population stability across multiple studies. Entries refer to evolutionary change relative to their respective ancestral controls.

The evolution of constancy stability is understood to depend upon either a decline in intrinsic population growth rate (*r*)or increase in equilibrium population size (*K*) (Dey *et al* 2012). Actually, while much discussion of the evolution of stability centres around changes in these familiar parameters of simple population growth models, the operative mechanism is through the effects on the return map of elevated realized growth rates at high density, spanning below and above equilibrium population size (see Fig. 1 in Dey *et al* 2012). Selection for adaptation to larval crowding at high food amounts in the LCUs did not lead to a substantial evolutionary decline in *r*, or increase in *K* (Fig 3 a, b, c, d, Table 3 a, b, c, d). What did evolve in the LCUs was an elevated realized population growth rate, roughly spanning a range of densities from medium to equilibrium population size (Fig. 4 b). We suspect that the fact that LCUs, unlike the MCUs (see Fig. 5 in Pandey and Joshi 2022), did not evolve higher realized population growth rates at densities above the equilibrium population size is the explanation for why LCUs evolved enhanced persistence but not constancy. Assessing this speculation will require theoretical study of how changes in density-specific realized population growth rates affect the shape of the return map.

In tandem with such studies, we also need to develop a conceptual framework for understanding how changes in different fitness components, and their sensitivity to density, results in changes in the density-specific realized population growth rates. *Drosophila* populations subjected to chronic larval crowding at different combinations of egg number and food amount show considerable variation in the underlying traits through which they evolve greater competitive ability (Table 7). However, there is as yet no clear conceptual link between changes in fitness-related traits and in density-specific realized population growth rates. For example, both the MCUs and LCUs have evolved greater competitive ability, but through different life history traits, as the MCUs evolved greater time efficiency of food-to-biomass conversion and the LCUs evolved higher feeding rate at larval stage (Table 7). A trait like greater time efficiency of food-to-biomass conversion could be increasing pre-adult survivorship under crowding in MCUs (see Pandey et al 2022) which could contribute to higher *K* and lower *α* in MCUs. In contrast, the evolution of higher feeding rate in the LCUs can lead to greater mortality in population dynamics assay as compared to the MCUs, which can explain stability has evolved differently in these two populations which have experienced different types of larval crowding.

The role of larval ecology in shaping constancy stability becomes more evident when we compare to the LCUs with the CU populations which had been selected at a similar combination of egg number and food volume as the LCUs. Similar to the LCUs, the CUs had not evolved constancy stability (Mueller *et al* 2000). While persistence stability evolved in the LCUs, persistence could not be calculated in CUs because no extinctions occured due to the large population sizes at which the CUs (and UUs, the controls) were maintained (Mueller *et al* 2000). Similar to the LCUs, the CUs did not evolve any differences in the surrogates of *r* and *K* (Mueller *et al* 2000) relative to the control populations. Further, both the LCUs and CUs evolved higher pre-adult survivorship and faster development at high density; although CUs had evolved increased tolerance to metabolic waste (Shiotsugu *et al* 1997, Borash *et al* 1998) while LCUs did not (Sarangi 2018) (Table 7).

The indirect comparison of LCUs and MCUs, via transformed response variables scaled by control population values, yielded no significant differences between the two selection regimes for any of the response variables other than constancy measured as CV in population size (Table 6). This pattern is slightly discordant with a qualitative comparison of the LCU versus MB, and MCU versus MB results, which suggests that LCUs differ from MCUs not just in CV of population size, but also in estimates of *r* and *K*, and the pattern of density-specific realized population growth rates. We suspect the reason these additional differences were not picked up in the analysis of transformed response variable is due to reduced statistical power in the latter, as a result of additional error being introduced during the scalin with mean control population values.

In summary, it is clear that the impact of density-dependent selection on the evolution of population stability attributes can be quite nuanced, and seems to depend on the egg number and food amount combination at which the selection for adaptation to larval crowding was experienced, thereby validating the speculations of Dey *et al* (2012). It is clear that density-dependent selection can affect the evolution of constancy and persistence in very context-specific manners, and that further theoretical and experimental studies linking changes in fitness components, and their sensitivity to density, to consequent changes in the pattern of density-specific realized population growth rates and return maps will go a long way in enhancing our understanding of these important phenomena linking population ecology and evolution.

## ACKNOWLEDGMENTS

We thank Pratyusha Banerjee, Ramesh Kokile and Ennya Anna Thomas for help in the population dynamics xperimentss, and Avani Mital, Manaswini Sarangi, Srikant Venkitachalam, N. Rajanna and Muniraju for assistance in the maintenance of selected and control populations. NP was supported by a doctoral fellowship from the Jawaharlal Nehru Centre for Advanced Scientific Research. This work was supported by a J.C. Bose National Fellowship (SERB, Government of India) to AJ and, in part, by AJ’s personal funds.

## Notes

### Competing Interest Statement

The authors have declared no competing interest.

